# Prevalence of chloroquine and antifolate drug resistance markers in *Plasmodium falciparum* parasites in Rwanda, 2018

**DOI:** 10.64898/2026.07.31.742040

**Authors:** Sara L. Cantoreggi, Aline Uwimana, Jean Damascene Niyonzima, Aimable Mbituyumuremyi, Christian Nsanzabana

## Abstract

**Background:** Antimalarial drug efficacy is threatened by the development of drug resistance. In Rwanda, high levels of resistance have been reported in the past, including mutations in the *Pfk13* gene associated with partial resistance to artemisinin, leading to delayed parasite clearance of Artemether-Lumefantrine (AL). Therefore, continuous molecular surveillance of drug resistance markers is required. Here, we report polymorphisms in the *Pfcrt, Pfdhps*, and *Pfdhfr* genes in samples collected in Rwanda in 2018.

**Methods:** Samples from three sites (Masaka, Rukara, and Bugarama) were collected during a therapeutic efficacy study (TES) conducted in 2018. Targeted amplification of resistance genes of interest was performed by PCR, and products were sequenced by Sanger sequencing.

**Results:** The K76**T** *Pfcrt* mutation was found in 10% of the samples, only in Bugarama. For *Pfdhps*, the A437**G**, K540**E** and A581**G** mutations were detected in 94%, 94% and 55% of the samples, respectively. For *Pfdhfr*, the N51**I**, S198**N**, C59**R** and I164**L** mutations were found in 99%, 99%, 88% and 16% of samples, respectively. The latter was only detected in Masaka and Rukara.

**Conclusions:** We confirm the slow and partial recovery of chloroquine susceptibility in the country. All previously reported *Pfk13* mutants were *Pfcrt* wild type, thus possibly selected by AL. The prevalence of *Pfdhps* and *Pfdhfr* mutants, including those with the highly resistant I164L mutation, is high and increasing, despite the absence of drug pressure. It is imperative to closely monitor resistance and efficacy of AL and other treatment options as part of the malaria surveillance program and resistance mitigation strategy in the country.

## BACKGROUND

Malaria affects millions of people every year, particularly children in Sub-Saharan Africa (1). Interventions aimed at its control and elimination have been in place for decades, including the deployment of highly effective therapies. Antimalarial drug efficacy is, however, threatened by the development of drug resistance, which to date has been detected for almost all antimalarial drugs (2, 3). This represents a major challenge for malaria control and elimination.

Rwanda has a history of high prevalence of antimalarial drug resistance. *P. falciparum* malaria cases were treated with chloroquine (CQ) until 2001 and with amodiaquine+sulfadoxine-pyrimethamine (AQ+SP) until 2006, when Artemisinin-based Combination Therapies (ACTs) were introduced (4, 5). SP was introduced in 2005 for intermittent preventive treatment in pregnancy (IPTp) but discontinued in 2008 (6). The shifts in treatment policy were always dictated by high levels of drug resistance, which were reported in the 1990s for CQ (6) and right after introduction for AQ and SP (7, 8). In 2021, partial resistance to artemisinin was reported in the country. In the context of a therapeutic efficacy study (TES) conducted in 2018, SNPs in the *P. falciparum kelch 13* (*Pfk13*) gene were detected and associated with delayed parasite clearance in patients treated with Artemether-Lumefantrine (AL) (9).

Resistance to CQ is associated with polymorphisms in the *P. falciparum chloroquine resistance transporter* (*Pfcrt*) gene, specifically to the K76**T** point mutation (10). Evidence from other African countries shows that when CQ drug pressure is lifted, wild-type parasites re-emerge (11, 12), likely due to their fitness advantage over mutants. Additionally, *Pfcrt* wild-type (K76, or CVMNK haplotype) parasites may be selected by AL, as they have reduced susceptibility to lumefantrine (13). Similar observations have been made for parasites carrying the N**F**D haplotype (N86-184**F**-D1246) in the *multidrug resistance 1* (*Pfmdr-1*) gene in multiple African countries (14-17). The prevalence of K76**T** mutations in different sites across Rwanda has been reported to decrease substantially in the last fifteen years. For instance, in Huye District (Southern Province) (Figure 1), prevalence decreased from 76% (N = 124) in 2010 to 26% (N = 202) in 2023 (18). In Rukara (Bugesera District, Eastern Province), a recent study reported prevalence as low as 6% (N = 82) in 2021 (19).

**Figure 1.**
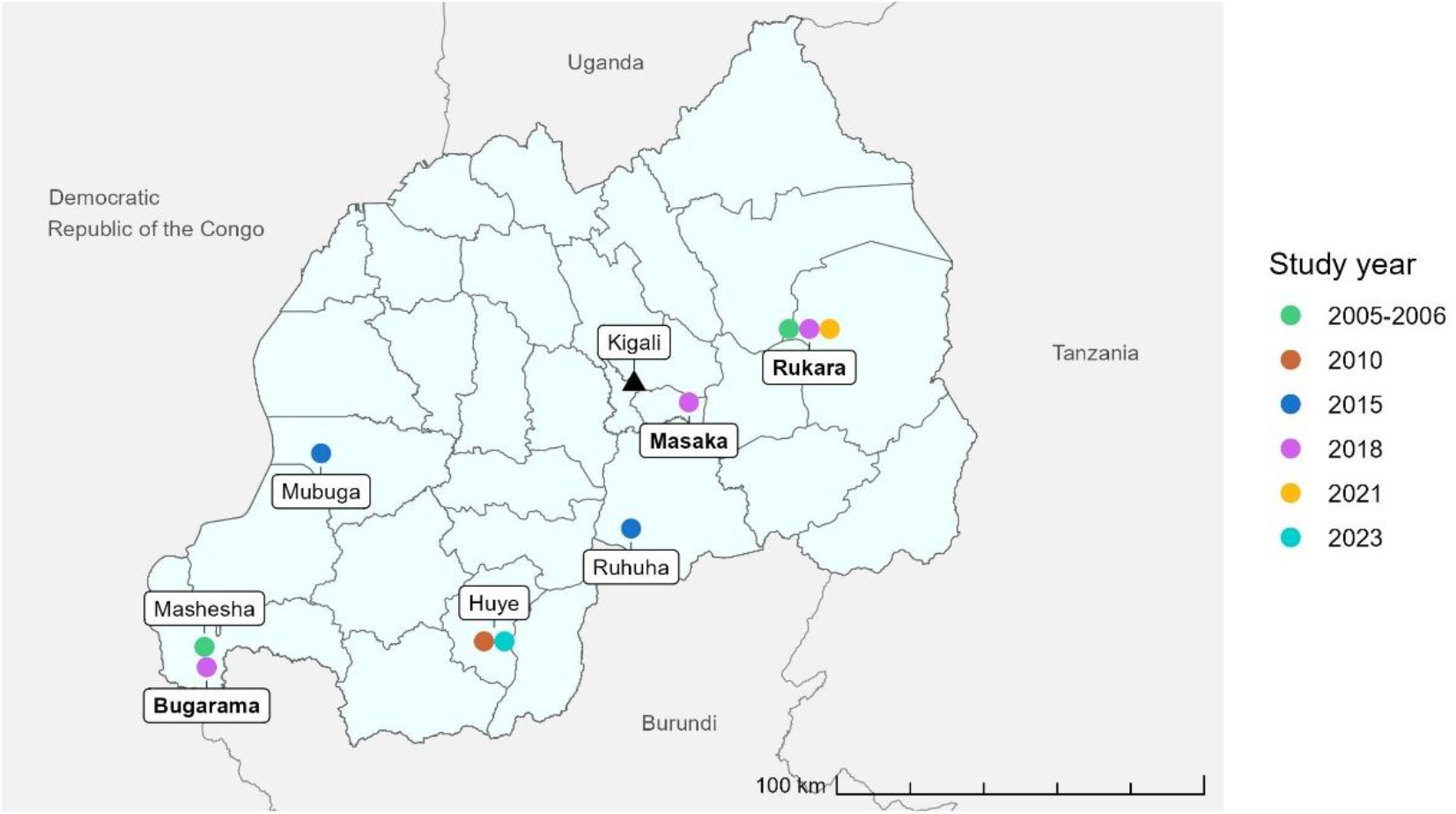
Map of Rwanda with study sites of current (bold) and other studies, colored by study year.

Resistance to SP is associated with multiple, combined SNPs in the *P. falciparum dihydropteroate synthase* (*Pfdhps*) and *dihydrofolate reductase* (*Pfdhfr*) genes (20-23). A *Pfdhps/Pfdhfr* quintuple mutation (*Pfdhps* double mutation (A437**G**+K540**E**) combined with *Pfdhfr* triple mutation (N51**I**+C59**R**+S108**N**)) increases the risk of SP resistance (24, 25), and the risk of treatment failure increases with increasing order of mutations (sextuple, septuple, or even octuple mutations). Sextuple mutations (*Pfdhps* A437**G**+K540**E**+A581**G** combined with *Pfdhfr* N51**I**+C59**R**+S108**N**) were reported at a prevalence ranging between 15% and 50% in different regions of the country in 2005, 2010 and 2015 (5, 8, 24). The highly pyrimethamine-resistant *Pfdhfr* I164**L** mutation (26) was reported at a prevalence of around 10% in 2005 (N = 725) and 2015 (N = 190), and then 25% (N = 77) in 2021, only in sites located in the Eastern Province (5, 8, 19).

Here, we report polymorphisms in the *Pfcrt, Pfdhps*, and *Pfdhfr* genes in samples collected in the 2018 TES conducted in Rwanda. Our data complements previously published data on *Pfk13* and *Pfmdr-1* polymorphisms (9), with the aim of obtaining a comprehensive drug resistance profile of *P. falciparum* in three sentinel sites in Rwanda in 2018.

## METHODS

### Sample collection

Samples were collected in a TES conducted in Rwanda in 2018. Samples were collected on dried blood spots (DBS on Whatman 903) in three sentinel sites: Masaka (Kicukiro District, City of Kigali), Rukara (Kayonza District, Eastern Province) and Bugarama (Rusizi District, Western Province) (Figure 1). Written informed consent was obtained from all study participants, and samples were anonymized and used only for study purposes. Detailed information about site characteristics and sample collection has been published elsewhere (9).

### Sample processing and sequencing

3 punches of 3 mm diameter were obtained from each DBS. DNA extraction was performed with the QIAamp 96 DNA Blood Kit (Qiagen) according to the manufacturer’s instructions (27). Genes *Pfcrt, Pfdhps*, and *Pfdhfr* were first amplified by primary and nested PCR, and then Sanger-sequenced on ABI3730XL (Thermo Fisher Scientific). Raw sequencing data was analyzed with SeqScape version 4 (Applied Biosystems) using the 3D7 strain as reference. The following SNPs were taken into consideration: M74**I**, N75**E**, K76**T** for *Pfcrt*; S436**A**, A437**G**, K540**E**, A581**G**, A613**S** for *Pfdhps*; N51**I**, C59**R**, S108**N**, I164**L** for *Pfdhfr*. Methods details are available in the Supplementary Files.

### Data analysis

Prevalence of SNPs among all isolates was calculated and then stratified by site in Excel 2016 (Microsoft). Mixed infections (mutant + wild type) were considered mutants. R studio (version 4.4.2) was used to compute 95% confidence intervals and produce all figures.

## RESULTS

256 samples (53 from Masaka, 103 from Rukara, and 100 from Bugarama) were successfully Sanger-sequenced for all genes.

### Pfcrt

The K76**T** SNP in *Pfcrt* was found in 10% (26/256) of the isolates. This was always detected together with the M74**I** and N75**E** SNPs, forming the CV**IET** haplotype. No other mutant haplotypes were detected. This haplotype was only found in Bugarama, where it had a prevalence of 26% (26/100), while all parasites found in Masaka and Rukara had the CVMNK (wild type) haplotype (Figure 2A). All *Pfk13* (R561**H** and P574**L**) mutants previously reported by Uwimana et al. were also found in these two sites (Figure 2B) (9), and all were CVMNK in *Pfcrt* (Figure 2D). Among them, 15 had the N**F**D, 12 the NYD (wild type) and 5 another (4 N**FY** and 1 NY**Y**) haplotype in *Pfmdr-1* (Figure 2D).

**Figure 2.**
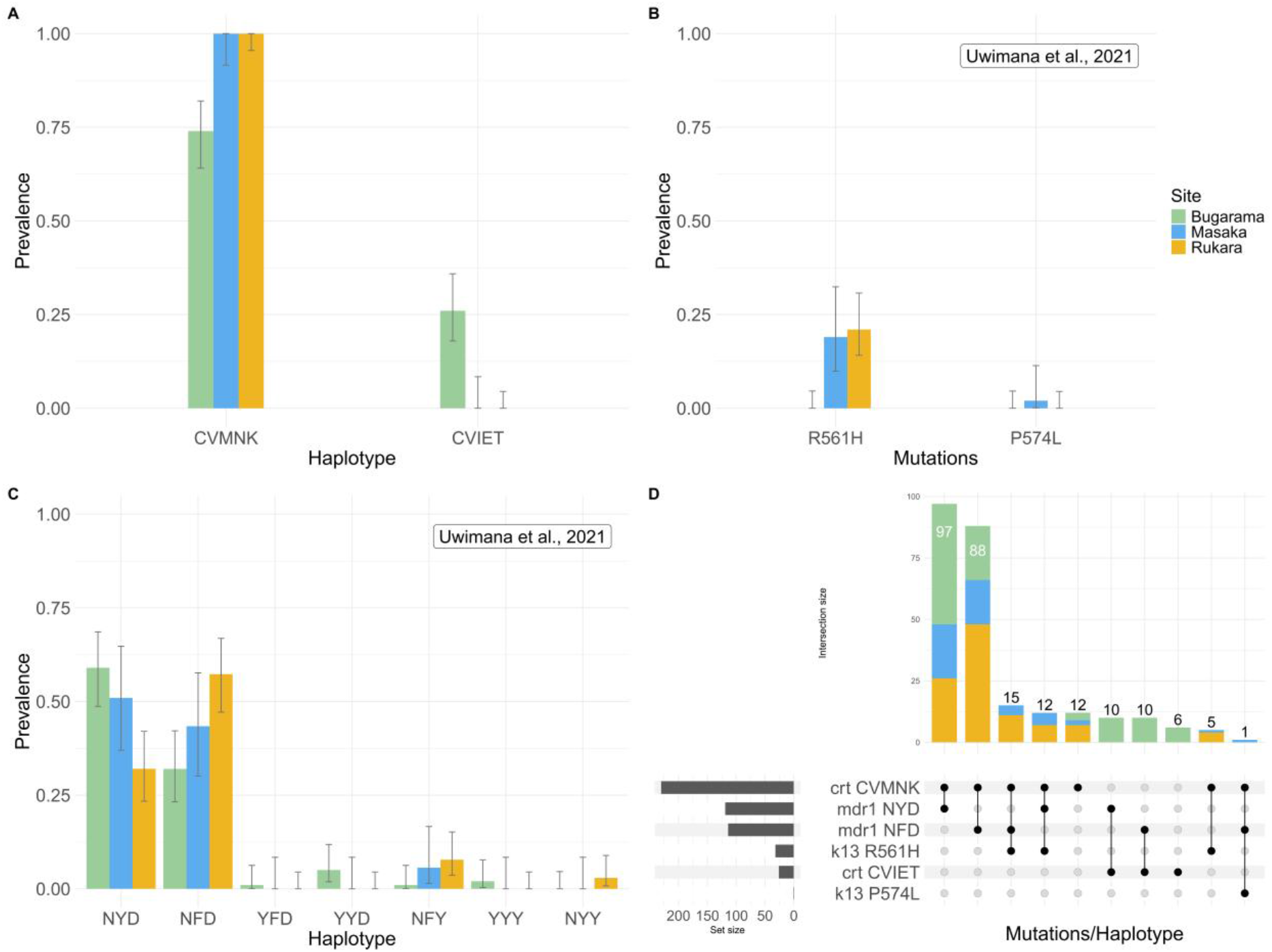
Prevalence of *Pfcrt* haplotypes (**A**), *Pfk13* mutations (**B**) and *Pfmdr-1* haplotypes (**C**) in Bugarama, Masaka, and Rukara. Data shown in B and C was published previously (9). Upset plot of *Pfk13* mutations, *Pfcrt* and *Pfmdr-1* haplotypes (**D**). Only selected haplotypes relevant for partial resistance to AL are shown.

### *Pfdhps* and *Pfdhfr*

*Pfdhps* and *Pfdhfr* mutations were detected at high prevalence in all three sites. For *Pfdhps*, the A437**G** and K540**E** SNPs were both found almost at fixation in 94% (240/256) of the samples, while the A581**G** SNP was detected in 55% (140/256) of the samples. No S436**A** or A613**S** SNPs were detected (Figure 3A). For *Pfdhfr*, the N51**I** and S198**N** SNPs were both at fixation in 99% (254/256) of the samples, while the C59**R** was found in 88% (226/256) of the samples. The I164**L** mutation was detected in 16% (42/256) of the samples, and only in Masaka (25%, 13/53) and Rukara (28%, 29/103) (Figure 3B). Sextuple (*Pfdhps* A437**G**+K540**E**+A581**G** and *Pfdhfr* N51**I**+C59**R**+S108**N**) mutations were found in 37% (95/256) of the samples. Quintuple (*Pfdhps* A437**G**+K540**E** and *Pfdhfr* N51**I**+C59**R**+S108**N**) and septuple (*Pfdhps* A437**G**+K540**E**+A581**G** and *Pfdhfr* N51**I**+C59**R**+S108**N**+I164**L**) mutations had lower prevalence of 30% (79/256) and 13% (34/256), respectively (Figure 3C/D).

**Figure 3.**
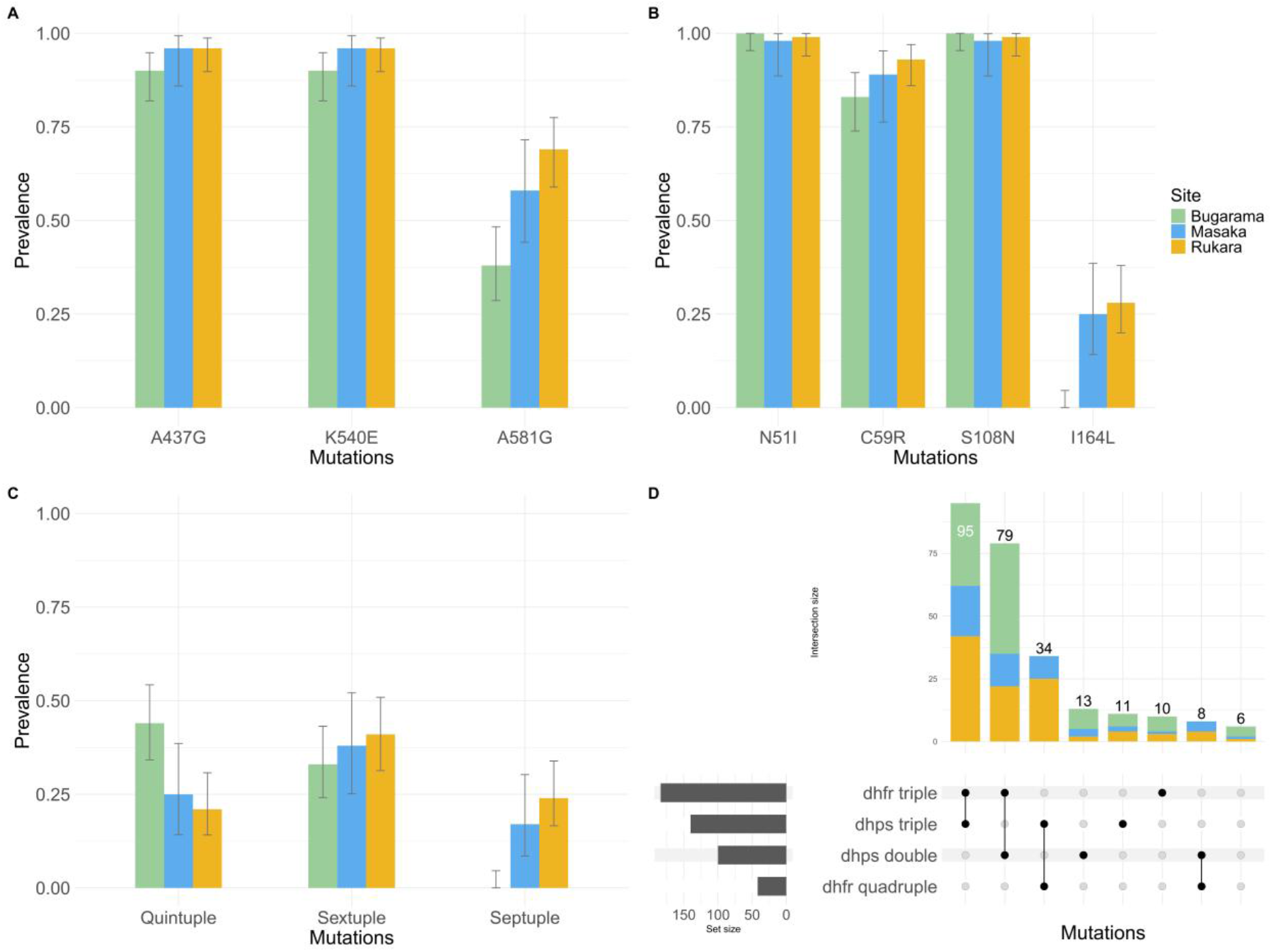
Prevalence of *Pfdhps* mutations (**A**), *Pfdhfr* mutations (**B**) and *Pfdhps*+*Pfdhfr* combinations (**C**) in Bugarama, Masaka, and Rukara. Upset plot of double and triple *Pfdhps* mutations and triple and quadruple *Pfdhfr* mutations (**D**).

## DISCUSSION

Molecular surveillance of drug resistance markers plays an important role in detecting and monitoring spatiotemporal trends of resistant parasites, thus providing useful information for the development of new strategies aimed at curbing the effects of drug resistance as well as the evaluation of additional antimalarials to be used in a country.

In Rwanda, chloroquine use was officially halted in 2001 after numerous reports of drug failure (6). The first molecular data available dates to 2010, when the prevalence of *Pfcrt* K76**T** mutants was 76% in Huye (18) (Figure 1). Prevalence was therefore maintained at high levels even ten years after discontinuation of chloroquine use, likely due to drug pressure given by AQ+SP until 2006 (24, 28). Thereafter, prevalence decreased and was reported to be 55% in Ruhuha (Bugesera District, Eastern Province) and 44% in Mubuga (Karongi District, Western Province) in 2015 (5). Only three years later, in 2018, our data shows a prevalence of 26% in Bugarama and 10% countrywide. Our findings confirm the slow and partial recovery of chloroquine-susceptibility in the country as reported previously (5), and align with the most recent reports from 2021 and 2023 (18, 19).

Interestingly, K76**T** (CV**IET**) mutant parasites were only detected in Bugarama, while none were detected in Masaka and Rukara, where *Pfk13* mutants have been reported (9). These results suggest that the use of AL in Rwanda may be selecting for *Pfk13* mutants (partially resistant to artemisinin) and wild type *Pfcrt* CVMNK parasites (less susceptible to lumefantrine). This hypothesis is further supported by the fact that about half of the *Pfk13* mutants also presented the N**F**D haplotype in the *Pfmdr-1* gene, and the prevalence of this haplotype across the country increased from 40% in 2010 (24) to 90% in 2018 (9). Multiple studies conducted in other African countries have reported selection of the CVMNK and N**F**D haplotypes by AL (13, 29, 30).

Despite SP officially not being in use in Rwanda since 2008, the prevalence of polymorphisms in the *Pfdhps* and *Pfdhfr* genes in the country is still very high. Indeed, we report *Pfdhps* double and *Pfdhfr* triple mutants to all have prevalence >80% in our sample set. 84% of the samples had at least a quintuple mutation and 50% at least a sextuple mutation. This aligns with previous studies, as high levels of these mutations were reported in Rukara and Mashesha (Rusizi District, Western Province) in 2005 (31), and Ruhuha and Mubuga in 2015 (5). We also report an increase of the highly resistant I164**L** SNP in *Pfdhfr*, with a prevalence of 28% in Rukara and 25% in Masaka in 2018. This is in line with recent reports from 2021 (19) and confirms the presence of this mutation only in sites located in the Eastern part of the country. Our data shows that, even ten years after cessation of SP use, the parasites circulating in the country continue to be for the most part mutants, and their prevalence, specifically that of the 164**L** mutation, is increasing. Three hypotheses could explain this, namely, (1) unauthorized use of SP, (2) selection by cotrimoxazole, and (3) parasite importation from neighboring countries.

1. SP may be a cheaper, and easier to administer alternative to ACTs and its misuse has been reported in several other African countries such as the Democratic Republic of the Congo (DRC), Tanzania, and Uganda (32); however, we believe it unlikely that in Rwanda SP would be misused to levels high enough to select for mutants.
2. Cotrimoxazole is commonly used in Rwanda for treatment of infections and as prophylaxis for opportunistic infections in People Living with HIV (PLHIV) (33), and may exert selective pressure on the parasites due to the chemical similarities between sulfadoxine and cotrimoxazole. Some studies have linked cotrimoxazole use to an increased risk of developing SP resistance (23, 34), while others have disproven this hypothesis (35-37). In our case, cotrimoxazole pressure would not explain the increase in I164**L** *Pfdhfr* mutations, which are linked to pyrimethamine rather than sulfadoxine.
3. As of 2018, all countries bordering Rwanda were using SP for IPTp (38), and some of them, including Uganda (39, 40) and Tanzania (41) have reported an increase of the I164**L** SNP in recent years. On the contrary, this SNP has so far been detected only in minimal prevalence in the DRC (42, 43) and in Burundi (44). Resistant parasites may therefore be imported from Uganda or Tanzania, which could also explain why the mutation was detected in Rukara and Masaka, both located on main travel axes between Rwanda, Uganda and Tanzania, and not in Bugarama, located at the border with the DRC and Burundi in the Western part of the country. We believe this hypothesis to be the most plausible, but additional population genomics studies are needed to confirm it.

Our study offers a valuable insight into the antimalarial resistance markers landscape in 2018 in Rwanda, but not without limitations. First, because this and previous studies were not always conducted in the same sentinel sites, it is difficult to make accurate comparisons of the results and analyze trends of resistant clones over time. Despite the moderate dimensions of the country, the Rwandan parasite population seems to be highly diverse (45), and a generalization of the drug resistance trends to the country level may therefore not be possible. Population genomics studies may help to further understand population structure and confirm if the parasites found in Rukara and Masaka are genetically distinct from those found in Bugarama, and if they do indeed share ancestry with parasites from neighboring countries. Additionally, due to the nature of Sanger sequencing, clones present in the samples in minor concentration (i.e., minority clones) are likely to be missed and thus not included in prevalence estimates.

In conclusion, our findings, combined with those obtained previously by Uwimana et al. (9), provide an overview of the drug resistance markers landscape in Rwanda in 2018. Besides the P*fk13* mutations reported previously, our findings, such as the presumed selection of *Pfcrt* wild type clones by lumefantrine and the very high and increasing prevalence of *Pfdhps* and *Pfdhfr* mutations, are a cause of concern. It is therefore imperative to continue to closely monitor the development of these mutations, as well as the efficacy of AL and other treatment options, as part of the malaria surveillance program in the country.

## ETHICS

Samples used for this study were collected during the 2018 TES, which was approved by the Rwanda National Ethics Committee (reference 195/RNEC/2017), Rwanda National Health Research Committee, and the Johns Hopkins Bloomberg School of Public Health Institutional Review Board. Written informed consent was obtained from parents or guardians of study participants as indicated in (9). Samples were anonymized and only used for study purposes. All procedures were carried out according to the relevant ethical regulations.

## DATA AVAILABILITY

Sanger sequencing data will be made available upon reasonable request.

## ACKNOWLEDGEMENTS

We would like to thank the Rwanda Biomedical Centre (Rwanda) for sharing the samples used for this study. We also thank the U.S. President’s Malaria Initiative, U.S. Centers for Disease Control and Prevention (Rwanda) for their support in running the 2018 TES.

## AUTHOR INFORMATION

### Authors and Affiliations

**Swiss Tropical and Public Health Institute, Allschwil, Switzerland**

Sara L. Cantoreggi, Aimable Mbituyumuremyi, Christian Nsanzabana

**University of Basel, Basel, Switzerland**

Sara L. Cantoreggi, Aimable Mbituyumuremyi, Christian Nsanzabana

**Malaria and Other Parasitic Diseases Division, Rwanda Biomedical Centre, Kigali, Rwanda**

Aline Uwimana, Jean Damascene Niyonzima, Aimable Mbituyumuremyi

### Contributions

S.L.C.: laboratory analysis, data analysis, visualization, writing (original draft). A.U., J.D.N, A.M.: sample procurement(2018 TES). C.N.: conceptualization, supervision, funding acquisition, resources, writing (review and editing). All authors reviewed and approved the final version of this manuscript.

