## Supplementary Files for "Prevalence of chloroquine and antifolate drug resistance markers in *Plasmodium falciparum* parasites in Rwanda, 2018"

**Supplementary Table 1**. PCR primers sequences. 3

**Supplementary Table 2**. Thermocycling conditions for primary PCRs of all markers. 3

**Supplementary Table 3**. Thermocycling conditions for nested PCRs of Pfdhps and Pfcrt. 3

**Supplementary Table 4**. Thermocycling conditions for nested PCRs of Pfdhfr. 4

**Supplementary Table 5**. Positive controls and SNP variants. 4

**Supplementary Table 6**. Sequencing primers. 4

### METHODS

PCR protocols were adapted from previously published protocols (31, 55). Primers and amplification conditions are indicated below.

**Supplementary Table 1**. PCR primers sequences.

| Marker | Primer name | Reaction | Sequence 5’-3’ |
|---|---|---|---|
| ***Pfcrt*** | crt_fw_P3 | Primary PCR | TGGCTCACGTTTAGGTGGAG |
| | crt_rv_P3 | Primary PCR | TTCCCTTTTTATTTCCAAATAAGGAATA |
| | CRT_N_fw | Semi-nested PCR | CTTGTCTTGGTAAATGTGCTC |
| | crt_rv_P3 | Semi-nested PCR | TTCCCTTTTTATTTCCAAATAAGGAATA |
| ***Pfdhps*** | DHPS_P_fw | Primary PCR | ATTTTTGTTGAACCTAAACGTGCTGTTCA |
| | DHPS_P_rv | Primary PCR | CTTGTCTTTCCTCATGTAATTCATCT |
| | DHPS_N_fw2 | Nested PCR | CTAAACGTGCTGTTCAAAGAATG |
| | DHPS_N_rv2 | Nested PCR | CAATTGTGTGATTTGTCCACAA |
| ***Pfdhfr*** | DHFR_P_fw | Primary PCR | tttATGATGGAACAAGTCTGC |
| | DHFR_P_rv | Primary PCR | TAGTATATACATCGCTAACAGAAAT |
| | DHFR_N_fw | Semi-nested PCR | ACAAGTCTGCGACGTTTTCGATATTTATG |
| | DHFR_N_rv | Semi-nested PCR | TAGTATATACATCGCTAACAGAAAT |

**Supplementary Table 2**. Thermocycling conditions for primary PCRs of all markers.

| Temperature | Time | Cycles |
|---|---|---|
| 95°C | 12:00 min | |
| 95°C | 00:30 min | |
| 52°C | 01:30 min | 30 |
| 72°C | 01:30 min | |
| 72°C | 05:00 min | |
| 4°C | ∞ | |

**Supplementary Table 3**. Thermocycling conditions for nested PCRs of *Pfdhps* and *Pfcrt*.

| Temperature | Time | Cycles |
|---|---|---|
| 95°C | 12:00 min | |
| 95°C | 00:30 min | |
| 58°C | 01:00 min | 30 |
| 72°C | 01:00 min | |
| 72°C | 05:00 min | |

| | |
|---|---|
| 4°C | ∞ |

**Supplementary Table 4**. Thermocycling conditions for nested PCRs of *Pfdhfr*.

| Temperature | Time | Cycles |
|---|---|---|
| 95°C | 12:00 min | |
| 95°C | 00:30 min | |
| 52°C | 01:30 min | 30 |
| 72°C | 01:30 min | |
| 72°C | 05:00 min | |
| 4°C | ∞ | |

**Supplementary Table 5**. Positive controls and SNP variants.

| Marker | SNP | 3D7 strain | Dd2 strain |
|---|---|---|---|
| ***Pfcrt*** | M74I | Wild-type | Mutant |
| | N75E | Wild-type | Mutant |
| | K76T | Wild-type | Mutant |
| ***Pfdhps*** | S436A | Wild-type | Wild-type |
| | A437G | Mutant | Mutant |
| | K540E | Wild-type | Wild-type |
| | A581G | Wild-type | Wild-type |
| | A613S | Wild-type | Wild-type |
| ***Pfdhfr*** | N51I | Wild-type | Mutant |
| | C59R | Wild-type | Mutant |
| | S108N | Wild-type | Mutant |
| | I164L | Wild-type | Wild-type |

**Supplementary Table 6**. Sequencing primers.

The same primers used for nested PCR (Tables S3 and S4) were used, except for Pfdhps, for which two additional primers were used.

| Marker | Primer name | Reaction | Sequence 5'-3' |
|---|---|---|---|
| ***Pfdhps*** | DHPS_N_fw2 | Nested PCR | CTAAACGTGCTGTTCAAAGAATG |
| | DHPS_N_rv2 | Nested PCR | CAATTGTGTGATTTGTCCACAA |
| | DHPS_1_fw | Sequencing | AAAAAAAAAAAACAAATTCTATAGTGTAG |
| | DHPS_2_rv | Sequencing | TTATAATTGGTTTCGCATCACA |
